# ZNF217-USP15 signaling loop regulates oncogenic phenotypes in ovarian cancer cells

**DOI:** 10.64898/2026.08.30.748158

**Authors:** Ayokunnumi Ogunsanya, Fatimah Alfaran, Swethakumar Basavarajaiah, Achuth Padmanabhan

**Affiliations:** Department of Biological Sciences, University of Maryland Baltimore County, Baltimore, USA; University of Maryland Greenbaum Comprehensive Cancer Center, Baltimore, USA

## Abstract

ZNF217 is an established oncogenic transcription factor that promotes cancer progression and therapeutic resistance; however, the mechanisms regulating ZNF217 protein abundance remain poorly understood. Here, we identify ubiquitin-specific peptidase 15 (USP15) as a critical regulator of ZNF217 stability and define a reciprocal USP15–ZNF217 signaling loop that sustains malignant phenotypes in ovarian cancer. Stable overexpression of ZNF217 in OVCA420 ovarian cancer cells enhanced proliferation, epithelial–mesenchymal transition, migration, invasion, and extracellular matrix adhesion. Notably, ZNF217 overexpression increased USP15 protein abundance without altering USP15 mRNA levels, whereas ZNF217 depletion reduced USP15 protein levels, suggesting post-transcriptional regulation. Conversely, USP15 depletion markedly reduced ZNF217 protein abundance while increasing ZNF217 mRNA levels, indicating that USP15 regulates ZNF217 predominantly at the post-transcriptional level. Proteasome inhibition restored ZNF217 protein levels following USP15 depletion, further demonstrating that USP15 promotes ZNF217 protein stability. Functionally, USP15 depletion in ZNF217-overexpressing ovarian cancer cells suppressed proliferation and multiple metastatic phenotypes, including migration, invasion, extracellular matrix adhesion, anoikis resistance, and multicellular aggregate formation. In vivo, USP15 depletion significantly reduced tumor progression and metastatic burden and prolonged survival in mice bearing ZNF217-driven ovarian tumors. Furthermore, USP15 depletion enhanced the sensitivity of ZNF217-overexpressing cells to carboplatin, paclitaxel, and doxorubicin. Collectively, these findings identify USP15 as an upstream regulator of ZNF217 protein stability and reveal a positive-feedback loop between USP15 and ZNF217 that reinforces oncogenic signaling. Targeting USP15 may therefore represent an indirect therapeutic strategy for suppressing ZNF217-driven ovarian cancer, particularly given the challenges associated with directly targeting oncogenic transcription factors.

## 1. Introduction

Ovarian cancer is an extremely lethal gynecologic malignancy and a leading cause of cancer-related deaths among women [1]. Due to the absence of reliable early diagnostic markers, more than 70% of patients are diagnosed with metastatic disease [2]. Current therapies are ineffective against advanced disease, and consequently, the 5-year survival rate remains below 30% [3]. Improving these outcomes requires better understanding of the molecular factors that drive ovarian cancer metastasis and the mechanisms that regulate their activity in tumor cells.

Zinc Finger Protein 217 (ZNF217) is a potent oncogene frequently overexpressed in multiple malignancies, including ovarian cancer [4-8]. Although ZNF217 promotes metastasis and chemoresistance, the mechanisms underlying these effects and those governing ZNF217 protein stability and abundance remain poorly defined [4]. Here, we identify the deubiquitinase USP15 as a critical effector of ZNF217 and, to our knowledge, the first identified regulator of ZNF217 protein stability. We demonstrate that USP15 sustains ZNF217 protein abundance and is required for ZNF217-driven oncogenic phenotypes in ovarian cancer. Importantly, USP15 depletion impairs ZNF217-driven tumorigenesis in vivo and sensitizes ovarian cancer cells to extant chemotherapeutics. These findings reveal a previously unrecognized mechanism regulating ZNF217 protein homeostasis and identify USP15 as a potential therapeutic target in ZNF217-driven tumors.

## 2. Materials and Methods

### 2.1 Cell Lines and Plasmids

TYK-Nu, OVCAR3, and OVCA420 cells were obtained from Dr. K.K Wong (MD Anderson Cancer Center, USA) and cultured in RPMI-1640 medium supplemented with 10% FBS at 37°C/5% CO_2_. pMD2.G (#12259) and psPAX2 (#12260) vectors were obtained from Addgene. USP15, ZNF217, and GFP shRNA were purchased from SigmaAldrich.

### 2.2 Cell Viability Assay

5 × 10^3^ cells were seeded in a 6-well plate and allowed to grow for 5-7 days. Cells were then fixed using 4% PFA and stained with crystal violet, imaged, and quantified using ImageJ. For WST-1 assays 5000 cells were seeded in a 96-well plate and cell viability was determined after 72-96 hours.

### 2.3 Transwell Migration Assay

3-5 × 10^4^ cells were seeded in RPMI containing 10% FBS inside 24-well inserts. RPMI containing 20% FBS was added to the 24-well dish, beneath the permeable insert. The cells were incubated for 24-48 hours at 37°C/5% CO_2_. Migratory cells were fixed using 4% PFA, stained using crystal violet, imaged, and quantified using ImageJ.

### 2.4 Matrigel Invasion Assay

24-well permeable inserts were coated with 40□ µl of Matrigel (1:3 ratio) and allowed to set at room temperature. Inserts were placed in a well in a 24-well plate containing RPMI media with 20% FBS. 0.5-1 × 10□ cells were reconstituted in RPMI media with 10% FBS, added to the inserts, and incubated at 37°C/5% CO_2_ for 24-72 hours. Cells that invaded through the Matrigel were fixed with 4% PFA, stained with crystal violet, imaged, and quantified using ImageJ.

### 2.5 In vivo Mouse Experiments

10^7^ cells stably expressing luciferase were injected intraperitoneally into 6-8 weeks-old female Foxn1 nude mice. Tumor progression was monitored using the IVIS system after injecting 150 mg/Kg D-luciferin intraperitoneally. Signal intensity was quantified by Living Image analysis software.

### 2.6 Quantification and Statistical Analysis

All in vitro experiments were repeated thrice, and data is presented as mean ± standard deviation. Statistical analyses were performed using GraphPad Prism 10.6.1. Comparisons between two groups were performed using Welch’s two-tailed t-test. In vivo data were analyzed by two-way ANOVA test. Mouse survival was assessed using Kaplan–Meier analysis and the log-rank (Mantel–Cox) test. Statistical significance was defined as p < 0.05, *p < 0.005, **p < 0.001, and ***p < 0.0001.

## 3. Results

### 3.1 ZNF217 promotes oncogenic properties in ovarian cancer cells

To validate the oncogenic functions of ZNF217 in ovarian cancer cells, we generated OVCA420 cells stably overexpressing ZNF217 (Fig. 1A). ZNF217 overexpression markedly increased cell viability and proliferation (Fig. 1B). ZNF217 overexpression resulted in reduced epithelial and increased mesenchymal marker expression, indicative of EMT (Fig. 1C). Consistent with these molecular changes OVCA420-ZN217 cells exhibited enhanced ability to migrate (Fig. 1D), invade through Matrigel (Fig. 1E), and attach to extracellular matrix proteins such as fibronectin (Fig. 1F). These findings confirm ZNF217’s ability to drive oncogenic phenotypes in ovarian cancer cells.

**Figure 1.**
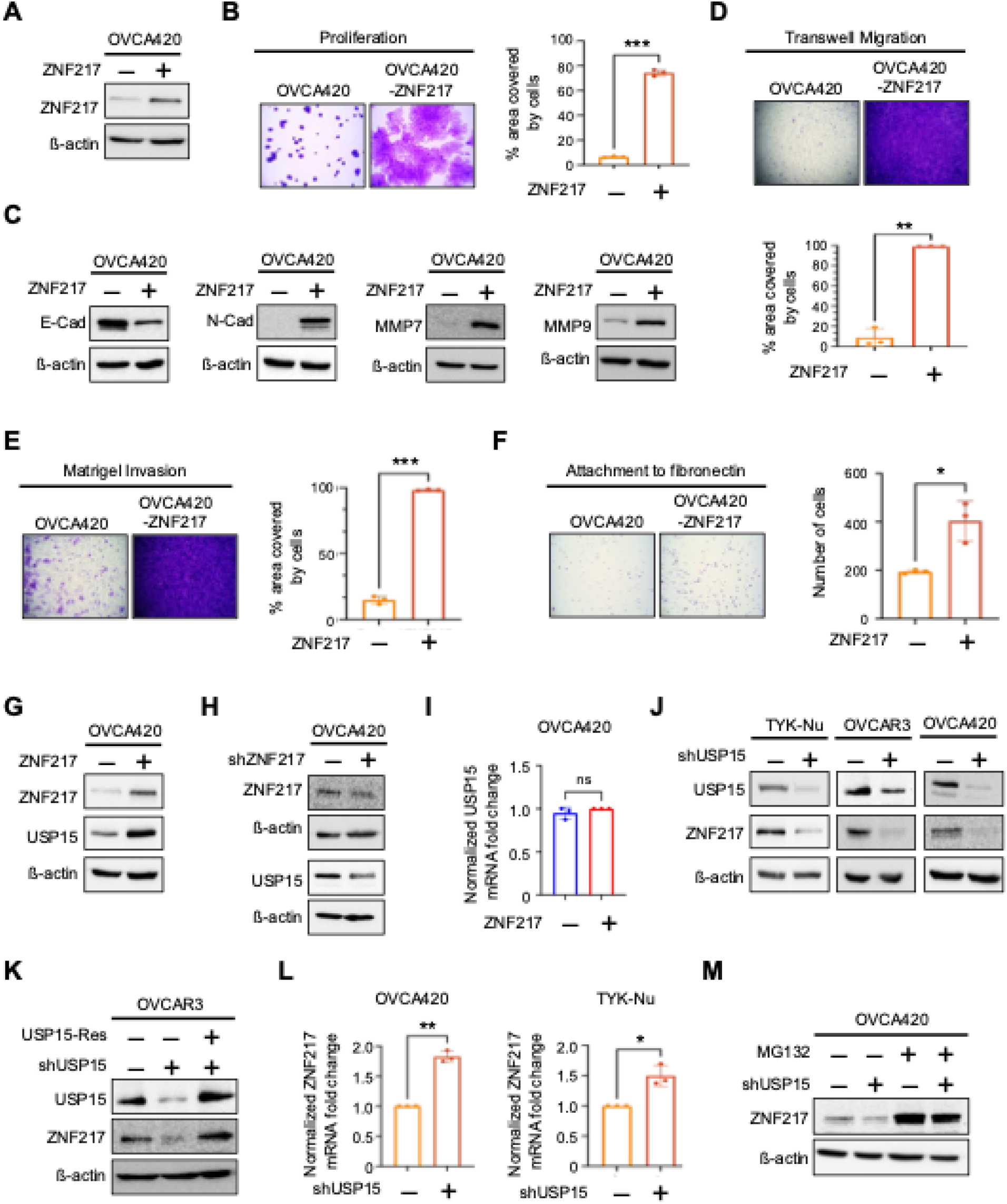
ZNF217 and USP15 reciprocally regulate each other’s levels in ovarian cancer cells. (A) Western blot confirming stable overexpression of ZNF217 in OVCA420 cells. (B) Representative crystal violet staining and quantified data shows that ZNF217 promotes cell viability and proliferation in OVCA420 cells. (C) ZNF217 overexpression causes a decrease in E-cadherin and increase in N-cadherin, MMP7 and MMP9 levels. (D) Transwell migration assay reveals an increase in migration upon ZNF217 overexpression in OVCA420 cells. (E) Matrigel invasion assay reveals that ZNF217 increases invasive potential in OVCA420 cells. (F) ZNF217 promotes attachment of OVCA420 cells to the extracellular matrix protein fibronectin. (G) Western blot analysis reveal that ZNF217 overexpression causes an increase in USP15 levels in OVCA420 cells. (H) shRNA-mediated ZNF217 knockdown in OVCA420 cells causes a decrease in USP15 levels. (I) RT-qPCR analysis show that ZNF217 overexpression does not alter USP15 mRNA levels (J) shRNA-mediated USP15 knockdown result in ZNF217 depletion in TYK-Nu, OVCAR3, and OVCA420 cells. (K) Ectopic expression of shRNA-resistant USP15 rescues ZNF217 levels in USP15 depleted OVCAR3 cells. (L) USP15 depletion causes an increase in ZNF217 levels in OVCA420 and TYK-Nu cells. (M) Proteasome inhibitor MG132 (10 µM, 6h) rescues USP15 knockdown induced depletion of ZNF217 in OVCA420 cells.

### 3.2 ZNF217-USP15 signaling loop regulates abundance of ZNF217 and USP15

ZNF217 overexpression in ovarian cancer cells resulted in an increase in USP15 protein levels (Fig. 1G). Conversely, ZNF217 knockdown depleted USP15 levels (Fig. 1H). RT-qPCR analysis revealed no change in USP15 mRNA level upon ZNF217 overexpression, suggesting that ZNF217 regulates USP15 abundance through post-transcriptional mechanisms (Fig. 1I). Interestingly, USP15 depletion markedly reduced ZNF217 levels, suggesting a reciprocal USP15–ZNF217 regulatory loop. (Fig. 1J). Ectopic expression of an shRNA-resistant USP15 in USP15-depleted OVCAR3 cells restored ZNF217 levels, confirming that USP15 regulates ZNF217 abundance (Fig. 1K). RT-qPCR analysis revealed that USP15 knockdown, in fact, increased ZNF217 mRNA levels, suggesting a discordance between ZNF217 mRNA and protein levels (Fig. 1L). These findings suggest that USP15 regulates ZNF217 predominantly through a post-transcriptional mechanism. Furthermore, proteasome inhibitor MG132 treatment restored ZNF217 protein levels in USP15-depleted OVCA20 cells, indicating that USP15 depletion promotes proteasome-dependent ZNF217 degradation (Fig. 1M). Collectively, these findings identify a reciprocal USP15–ZNF217 regulatory loop and establish USP15 as a critical regulator of ZNF217 protein stability. To our knowledge, this is the first report identifying an upstream regulator of ZNF217 stability in any cellular context.

### 3.3 USP15 depletion reverses ZNF217’s oncogenic potential in ovarian cancer cells

As USP15 depletion reduced ZNF217 protein levels, we next evaluated USP15’s utility as a therapeutic target in ZNF217-overexpressing ovarian cancer cells by stably depleting USP15 using a validated shRNA in OVCA420-ZNF217 cells (Fig. 2A). OVCA420-ZNF217 cells expressing GFP-targeting shRNA served as control (Fig. 2A). USP15 depletion significantly reduced cell viability and proliferation in both the short term (72 hours; Fig. 2B) and long term (7 days; Fig. 2C), as assessed by WST-1 assay and crystal violet staining, respectively.

**Figure 2.**
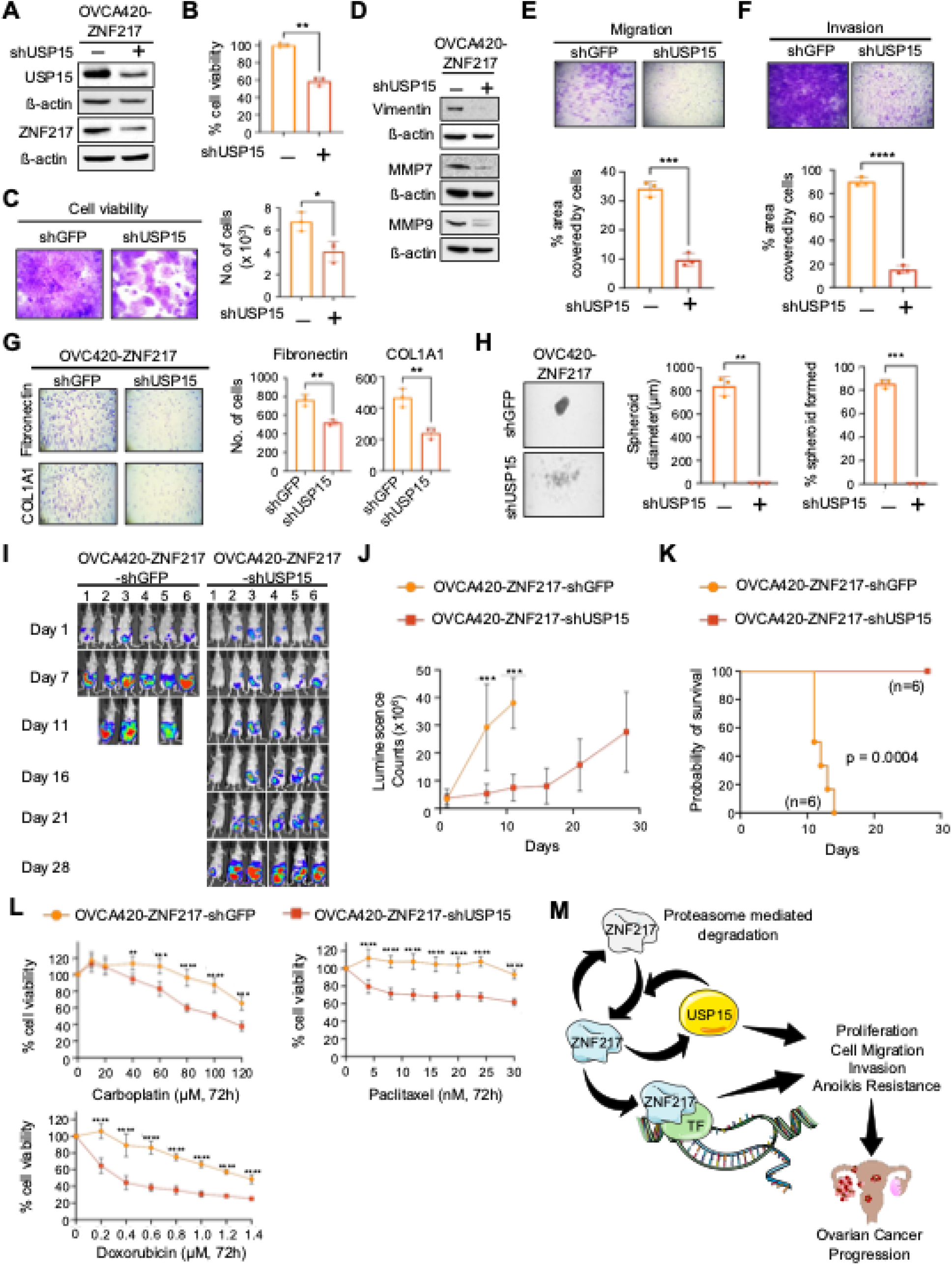
USP15 depletion impairs ZNF217 driven ovarian tumor progression and sensitizes ovarian cancer cells to chemotherapeutic drugs. (A) Western blot confirming stable USP15 knockdown in OVCA420-ZNF217 cells. USP15 depletion causes a decrease in ZNF217 level in these cells. (B) WST-1 assay show that USP15 depletion significantly decreases the viability of OVCA420-ZNF217 cells (96 hours). (C) Crystal violet staining show that USP15 depletion significantly decreases the viability and proliferation of OVCA420-ZNF217 cells (5 days). (D) USP15 depletion decreases the level of mesenchymal marker vimentin and ECM modifying enzymes MMP7 and MMP9 in OVCA420-ZNF217 cells. (E) Transwell assay show that USP15 depletion decreases cell migration in OVCA420-ZNF217 cells. (F) Matrigel invasion assay show that USP15 depletion decreases invasive potential of OVCA420-ZNF217 cells. (G) USP15 knockdown impairs the ability of OVCA420-ZNF217 cells to attach to extracellular matrix proteins such as fibronectin and collagen. (H) USP15 knockdown inhibits the ability of OVCA420-ZNF217 cells to form multicellular aggregates (spheroids) under low-attachment conditions. (I) Non-invasive imaging show decreased bioluminescent signal (tumor burden) in Foxn1 nude mice injected with OVCA420-ZNF217-shUSP15 cells compared to OVCA420-ZNF217-shGFP (control) cells. (J) Quantified bioluminescent signal from Foxn1 nude mice intraperitoneally injected with luciferase-tagged OVCA420-ZNF217-shGFP and OVCA420-ZNF217-shUSP15 cells show significant reduction in tumor burden upon USP15 depletion. N = 6 mice/group. (K) Foxn1 nude mice bearing OVCA420-ZNF217-shUSP15 tumors survive significantly longer than mice bearing OVCA420-ZNF217-shGFP tumors. (L) WST-1 assay shows that USP15 depletion in OVCA420-ZNF217 cells sensitizes them to carboplatin, paclitaxel, and doxorubicin (n=3, 72 hours). (M) Model summarizing ZNF217-USP15 regulatory loop in ovarian cancer cells and its impact on ovarian tumor progression and chemoresistance.

Furthermore, USP15 knockdown reduced expression of the mesenchymal marker vimentin as well as extracellular matrix-remodeling enzymes such as MMP7 and MMP9, suggesting attenuation of metastatic phenotypes (Fig. 2D). Consistent with these molecular changes, USP15 knockdown impaired metastatic phenotypes such as cell migration (Fig. 2E), invasion (Fig. 2F), adhesion to extracellular matrix proteins fibronectin and collagen (Fig. 2G), and resistance to anoikis and formation of multicellular aggregates/spheroids (Fig. 2H). Collectively, these findings demonstrate that USP15 is required for the maintenance of ZNF217-driven malignant phenotypes and identify USP15 as a potential therapeutic vulnerability in ZNF217-overexpressing ovarian cancer.

### 3.4 USP15 depletion inhibits ZNF217-driven tumor progression and sensitizes ovarian cancer cells to chemotherapy

To extend these findings to an in vivo model, we intraperitoneally injected luciferase-tagged OVCA420-ZNF217-shGFP and OVCA420-ZNF217-shUSP15 cells into 6–8-weeks-old female Foxn1 nude mice. USP15 depletion significantly reduced tumor progression and metastatic tumor burden in vivo (Figs. 2I and 2J) and prolonged survival in tumor-bearing mice (Fig. 2K). Further, consistent with ZNF217’s ability to promote chemoresistance in ovarian cancer cells, USP15 knockdown sensitized OVCA420-ZNF217 cells to carboplatin, paclitaxel, and doxorubicin (Fig. 2L). Taken together, these data support USP15’s role a critical mediator of ZNF217-driven tumor progression and highlights its potential as a therapeutic target in ZNF217-driven ovarian cancer.

## 4. Discussion and Conclusion

Although ZNF217 is an established oncogene that promotes tumor progression and therapeutic resistance, mechanisms regulating its protein abundance remain poorly understood [4-8]. Our data identify a signaling loop involving USP15 and ZNF217 that regulates each other’s abundance in cells (Fig. 2M). The reciprocal regulation of USP15 and ZNF217 suggests a positive-feedback loop that reinforces ZNF217 abundance and oncogenic activity (Fig. 2M). Consistent with this model, USP15 depletion suppressed proliferation and multiple ZNF217-driven metastatic phenotypes in vitro and in vivo. These findings identify USP15 as a critical mediator of ZNF217-driven tumorigenesis and suggest that targeting USP15 may provide an indirect strategy to suppress tumors driven by elevated ZNF217, an attractive possibility given the challenges of directly targeting transcription factors.

USP15 is overexpressed in several cancers and promotes tumor progression by stabilizing oncogenic signaling proteins and impacting multiple key processes [9-13]. Our findings extend these observations by identifying ZNF217 as a previously unrecognized component of the USP15-regulated proteostatic network. Future studies determining whether ZNF217 is a direct USP15 substrate and evaluating selective USP15 inhibition will be important for establishing the therapeutic potential of targeting the USP15–ZNF217 axis in ovarian cancer. In summary, we identify USP15 as an upstream regulator of ZNF217 protein stability and define a positive-feedback mechanism that sustains ZNF217-driven malignant phenotypes. Targeting this regulatory axis may represent a promising therapeutic strategy for ZNF217-driven ovarian tumors.

## Ethics Statement

Experiments in mice were performed as per a protocol (#1947) approved by the IACUC at UMBC.

## CRediT authorship contribution statement

Ayokunnumi Ogunsanya: Investigation, Methodology, Resources, Formal analysis, writing - review & editing. Fatimah Alfaran: Investigation, Formal analysis, writing - review & editing. Swethakumar Basavarajaiah: Investigation, writing - review & editing. Achuth Padmanabhan: Conceptualization, Formal analysis, Funding acquisition, Supervision, Writing - original draft, Writing - review & editing.

